# LncRNA 3222401L13Rik Is Up-regulated in Aging Astrocytes and Regulates Neuronal Support Function Through Interaction with Npas3

**DOI:** 10.1101/2024.08.14.607966

**Authors:** Sophie Schroeder, M. Sadman Sakib, Dennis Krüger, Tonatiuh Pena, Susanne Burkhardt, Anna-Lena Schütz, Farahnaz Sananbenesi, André Fischer

## Abstract

Aging is linked to a decline in cognitive functions and significantly increases the risk of neurodegenerative diseases. While molecular changes in all central nervous system (CNS) cell types contribute to aging-related cognitive decline, the mechanisms driving disease development or offering protection remain poorly understood. Long non-coding RNAs (lncRNAs) have emerged as key regulators of cellular functions and gene expression, yet their roles in aging, particularly within glial cells, are not well characterized. In this study, we investigated lncRNA expression profiles in non-neuronal cells from aged mice. We identified 3222401L13Rik, a previously unstudied lncRNA enriched in glial cells, as being specifically upregulated in astrocytes during aging. Knockdown of 3222401L13Rik in primary astrocytes revealed its critical role in regulating genes essential for neuronal support and synapse organization. This function was also conserved in human iPSC-derived astrocytes. Additionally, we found that 3222401L13Rik mediates its cellular effects through interaction with the transcription factor Neuronal PAS Domain Protein 3 (Npas3), and that overexpression of Npas3 effectively rescued the functional deficits observed in astrocytes lacking 3222401L13Rik. Our findings suggest that upregulation of 3222401L13Rik in aging astrocytes acts as a compensatory mechanism to enhance neuronal and synaptic support, potentially delaying the onset of molecular and structural changes in both astrocytes and neurons. Strategies to boost 3222401L13Rik expression earlier in life may help mitigate age-associated loss of neuronal plasticity.

## Introduction

Aging is a major risk factor for the onset and progression of neurodegenerative diseases (Hou *et al*, 2019). Even in the absence of pathological conditions, aging is associated with a decline in cognitive functions in both mice and humans (Berchtold *et al*, 2008) (Peleg *et al*, 2010) (Royall *et al*, 2005) (Stern, 2012} (Murman, 2015). Structural alterations in the aging brain include decreases in neuronal spine length and density, reduced memory-related neuronal firing, and changes in glial cell function, including altered astrocyte (Rodríguez *et al*, 2014) (Wang *et al*, 2011) (E., 2011) and microglia function (Hayakawa *et al*, 2007) (Hong *et al*, 2016). At the molecular level, shifts in gene expression have been described, reflecting processes such as synaptic transmission, vesicular transport, energy production, and immune activation (Peleg *et al*., 2010) (Lu *et al*, 2004) (Berchtold *et al*., 2008) (Stilling *et al*, 2014). However, the mechanisms underlying these changes in gene expression are still not fully understood, particularly concerning glial cells, when compared to neurons.

The non-coding RNAome has emerged as a critical regulator of gene expression and other cellular functions (Mattick *et al*, 2023), offering novel and so far unexplored opportunities for therapeutic interventions (Damase *et al*, 2021). Among non-coding RNAs, long non-coding RNAs (lncRNAs) represent a heterogeneous class of RNAs longer than 300 nucleotides that lack coding potential (Kapranov *et al*, 2007) (Mattick *et al*., 2023). Key characteristics of lncRNAs, in comparison to protein-coding RNAs, include their relatively lower abundance, preferential nuclear localization, greater tissue- and cell-specific expression patterns, and comparatively lower sequence conservation across species (Khorkova *et al*, 2015). Due to these features, lncRNAs were long considered as transcriptional noise, with research and drug discovery focusing primarily on protein-coding transcripts (Warner *et al*, 2018) (Liau *et al*, 2020). However, recent research has highlighted the significant roles of lncRNAs in physiological and pathological conditions, impacting a wide array of biological processes, including genome architecture regulation, gene expression modulation, splicing regulation, and control of protein translation and localization (Mattick *et al*., 2023). Additionally, the observation that approximately 40% of human lncRNAs exhibit brain-specific expression patterns suggests that these molecules play complex roles in brain development and function (Derrien *et al*, 2012), as well as in the pathogenesis of central nervous system disorders (Briggs *et al*, 2015) (Liau *et al*., 2020) (Musgrove *et al*, 2024). Several studies have demonstrated that lncRNAs are critical in cognitive diseases (Schröder *et al*, 2024) (Faghihi *et al*, 2008) (He *et al*, 2021) (Irwin *et al*, 2023) (Magistri *et al*, 2015) (Zhang *et al*, 2019) (Liau *et al*, 2023).

Given the limited research on the role of lncRNAs in the aging brain, and the sparse data on non-neuronal glial cells, this study aimed to identify lncRNAs differentially expressed in the hippocampus of aged versus young mice and to investigate the functional implications of candidate lncRNAs. Our findings reveal that one such deregulated lncRNA, *3222401L13Rik*, which has not been previously studied, is specifically upregulated in astrocytes of aged mice. Antisense oligonucleotide-mediated knockdown of *3222401L13Rik* in primary astrocytes, followed by total RNA sequencing, demonstrates its crucial role in regulating genes essential for neuronal support functions. This regulation impacts astrocytic processes such as glutamate uptake and calcium signaling, leading to disruptions in neuronal network plasticity and synapse density. Additionally, we show that its human homolog, *ENSG00000272070*, regulates similar genes and functions in human iPSC-derived astrocytes. Furthermore, we demonstrate that *3222401L13Rik* exerts its effects by interacting with and regulating the transcription factor Neuronal PAS Domain Protein 3 (Npas3). Importantly, overexpression of *Npas3* is sufficient to rescue the functional deficits observed in astrocytes lacking *3222401L13Rik*.

## Results

### Total RNA sequencing of glia of young and aged mice identifies differentially regulated lncRNAs

To identify deregulated lncRNAs in glia during aging, we isolated tissue from the hippocampal CA1 region of young (3 months) and old (16 months) mice. The hippocampus is a brain region crucial for memory function, and aging significantly affects this area. At 16 months, cognitive decline becomes observable in mice (Peleg *et al*., 2010). We processed the tissue to isolate nuclei via fluorescence-associated nuclei sorting (FANS), using neuronal nuclear protein (NeuN) to distinguish between neuronal (NeuN+) and non-neuronal nuclei (NeuN-) fractions. These two populations were then subjected to total RNA sequencing **(Fig. 1A)**.

**Figure 1:**
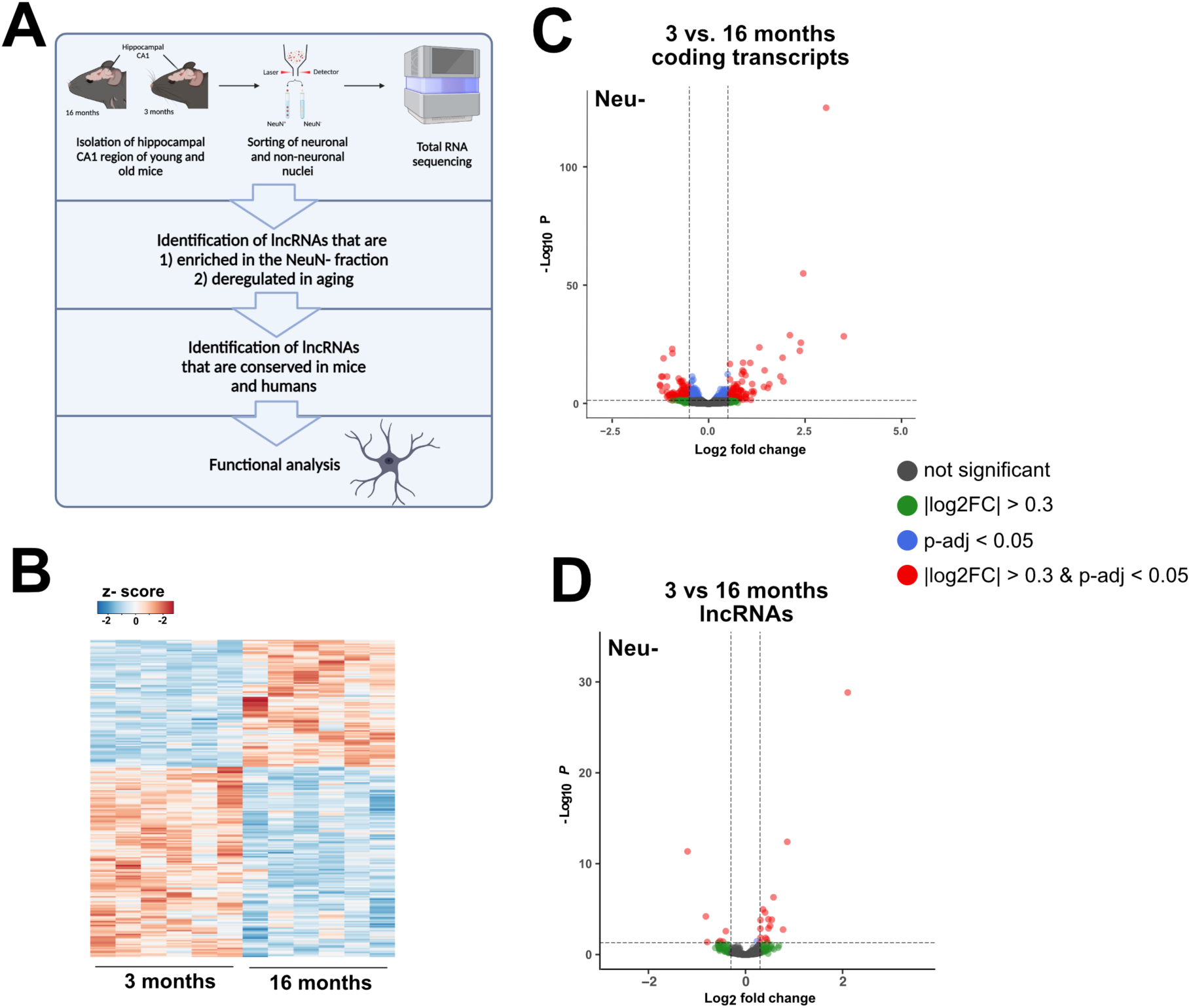
Aging induces changes in glial lncRNA expression patterns. **(A)** Schematic illustration of the experimental approach of the study. **(B)** Heatmap showing gene expression changes in Neu-nuclei isolated in 3 vs. 16 months mice. **(C)** Volcano plot showing the up- and downregulated coding transcripts when comparing Neu-nuclei from 3 vs 16-months-old mice **(D)** Volcano plot showing expression changes of lncRNAs in Neu-nuclei.

We aimed to identify lncRNAs that are enriched in the NeuN-fraction and deregulated during aging **(Fig. 1A)**. To achieve this, we first analyzed differentially regulated genes in the NeuN-populations of young and aged mice **(Fig. 1B, C)**. In comparison to young mice, 187 genes were upregulated and 310 genes were downregulated in 16-month-old mice (|log2FC| > 0.3, FDR < 0.05, basemean > 50) **(Table S1)**. Filtering for lncRNAs from this list, we identified 17 upregulated and 7 downregulated lncRNAs **(Fig. 1D)**. Although our analysis focused on NeuN-nuclei, the 17 deregulated lncRNAs may also be expressed in NeuN+ nuclei. To determine if any of these lncRNAs were particularly enriched in the NeuN-population, we compared gene expression profiles of the NeuN+ and NeuN-fractions. Substantial differences were observed, with 51 lncRNAs highly enriched in NeuN-nuclei **(Table S2)**.

Next, we compared the list of lncRNAs deregulated during aging (n = 17) with those enriched in the NeuN-fraction (n = 51). To facilitate the translation of our findings to human studies, we focused on lncRNAs with human homologs. Among the 17 upregulated lncRNAs in 16-month-old mice, seven were significantly enriched in the NeuN-fraction compared to NeuN+ nuclei (log2FC > 4, padj < 0.05), with only three having human homologs. These included nuclear enriched abundant transcript 1 (*Neat1*) and *3222401L13Rik*, both upregulated in 16-month-old mice, and *C430049B03Rik*, which was downregulated **(Table 1)**. Since *Neat1* has been extensively studied, particularly concerning its role in astrocytes (Irwin *et al*., 2023), and *C430049B03Rik* has been analyzed as a host gene for miR-322, miR-503, and miR-531 in mouse embryonic fibroblasts (Wang *et al*, 2019), we chose to focus our study on *3222401L13Rik*, a lncRNA that has not been previously characterized in any organ.

**Table 1:**
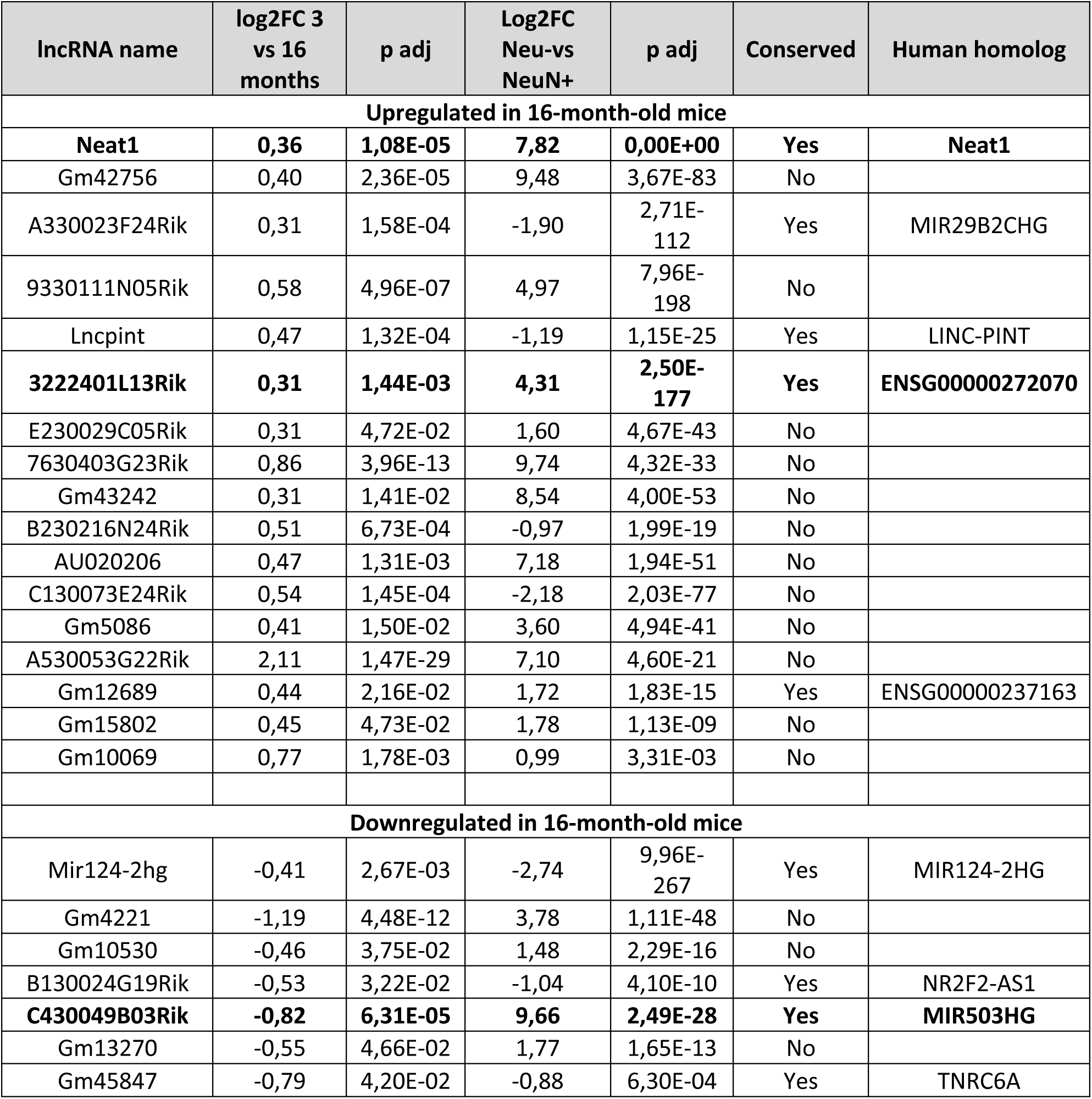
Characteristics of lncRNAs de-regulated during aging in Neu-nuclei.

| lncRNA name | log2FC 3 vs 16 months | p adj | Log2FC Neu-vs NeuN+ | p adj | Conserved | Human homolog |
| --- | --- | --- | --- | --- | --- | --- |
| **Upregulated in 16-month-old mice** |  |  |  |  |  |  |
| **Neat1** | **0,36** | **1,08E-05** | **7,82** | **0,00E+00** | **Yes** | **Neat1** |
| Gm42756 | 0,40 | 2,36E-05 | 9,48 | 3,67E-83 | No |  |
| A330023F24Rik | 0,31 | 1,58E-04 | -1,90 | 2,71E-112 | Yes | MIR29B2CHG |
| 9330111N05Rik | 0,58 | 4,96E-07 | 4,97 | 7,96E-198 | No |  |
| Lncpint | 0,47 | 1,32E-04 | -1,19 | 1,15E-25 | Yes | LINC-PINT |
| **3222401L13Rik** | **0,31** | **1,44E-03** | **4,31** | **2,50E-177** | **Yes** | **ENSG00000272070** |
| E230029C05Rik | 0,31 | 4,72E-02 | 1,60 | 4,67E-43 | No |  |
| 7630403G23Rik | 0,86 | 3,96E-13 | 9,74 | 4,32E-33 | No |  |
| Gm43242 | 0,31 | 1,41E-02 | 8,54 | 4,00E-53 | No |  |
| B230216N24Rik | 0,51 | 6,73E-04 | -0,97 | 1,99E-19 | No |  |
| AU020206 | 0,47 | 1,31E-03 | 7,18 | 1,94E-51 | No |  |
| C130073E24Rik | 0,54 | 1,45E-04 | -2,18 | 2,03E-77 | No |  |
| Gm5086 | 0,41 | 1,50E-02 | 3,60 | 4,94E-41 | No |  |
| A530053G22Rik | 2,11 | 1,47E-29 | 7,10 | 4,60E-21 | No |  |
| Gm12689 | 0,44 | 2,16E-02 | 1,72 | 1,83E-15 | Yes | ENSG00000237163 |
| Gm15802 | 0,45 | 4,73E-02 | 1,78 | 1,13E-09 | No |  |
| Gm10069 | 0,77 | 1,78E-03 | 0,99 | 3,31E-03 | No |  |
| **Downregulated in 16-month-old mice** |  |  |  |  |  |  |
| Mir124-2hg | -0,41 | 2,67E-03 | -2,74 | 9,96E-267 | Yes | MIR124-2HG |
| Gm4221 | -1,19 | 4,48E-12 | 3,78 | 1,11E-48 | No |  |
| Gm10530 | -0,46 | 3,75E-02 | 1,48 | 2,29E-16 | No |  |
| B130024G19Rik | -0,53 | 3,22E-02 | -1,04 | 4,10E-10 | Yes | NR2F2-AS1 |
| **C430049B03Rik** | **-0,82** | **6,31E-05** | **9,66** | **2,49E-28** | **Yes** | **MIR503HG** |
| Gm13270 | -0,55 | 4,66E-02 | 1,77 | 1,65E-13 | No |  |
| Gm45847 | -0,79 | 4,20E-02 | -0,88 | 6,30E-04 | Yes | TNRC6A |

The table displays the 17 lncRNAs that are significantly altered when comparing NeuN-nuclei from 3-month-old versus 16-month-old mice. It includes the corresponding fold change (log2FC 3 vs 16 months) and adjusted p-value (p adj). The column “log2FC Neu- vs Neu+” is based on differential expression analysis comparing transcripts in NeuN-versus NeuN+ nuclei, with a positive value above 4 indicating significant enrichment in NeuN-nuclei. The columns “Conserved” and “Human Homolog” indicate whether a human homolog exists.

### *3222401L13Rik* is a glial-enriched lncRNA showing increased expression in astrocytes of aged mice

*3222401L13Rik* is a long intergenic lncRNA (lincRNA) located on chromosome 18 in the mouse genome, with its human homolog (ENSG00000272070) found on chromosome 5 **(Fig. 2A)**. We confirmed the total RNA sequencing results with qPCR, demonstrating that 3222401L13Rik is predominantly expressed in NeuN-compared to NeuN+ nuclei **(Fig. 2B)** and exhibits increased expression in 16-month-old mice **(Fig. 2C)**. Since the NeuN-fraction includes all brain cell types except neurons, we further investigated the expression pattern of *3222401L13Rik* in glial cells by performing Magnetic Activated Cell Sorting (MACS) of astrocytes, oligodendrocytes, and microglia from 3-month-old mice. qPCR analysis showed that *3222401L13Rik* expression was comparable across all three glial cell types analyzed **(Fig. 2D)**. However, when comparing expression levels in 3-month-old and 16-month-old mice, we observed an increase in *3222401L13Rik* levels only in astrocytes, but not in oligodendrocytes or microglia **(Fig. 2E)**. Consequently, we focused on characterizing the role of 3222401L13Rik specifically in astrocytes.

**Figure 2:**
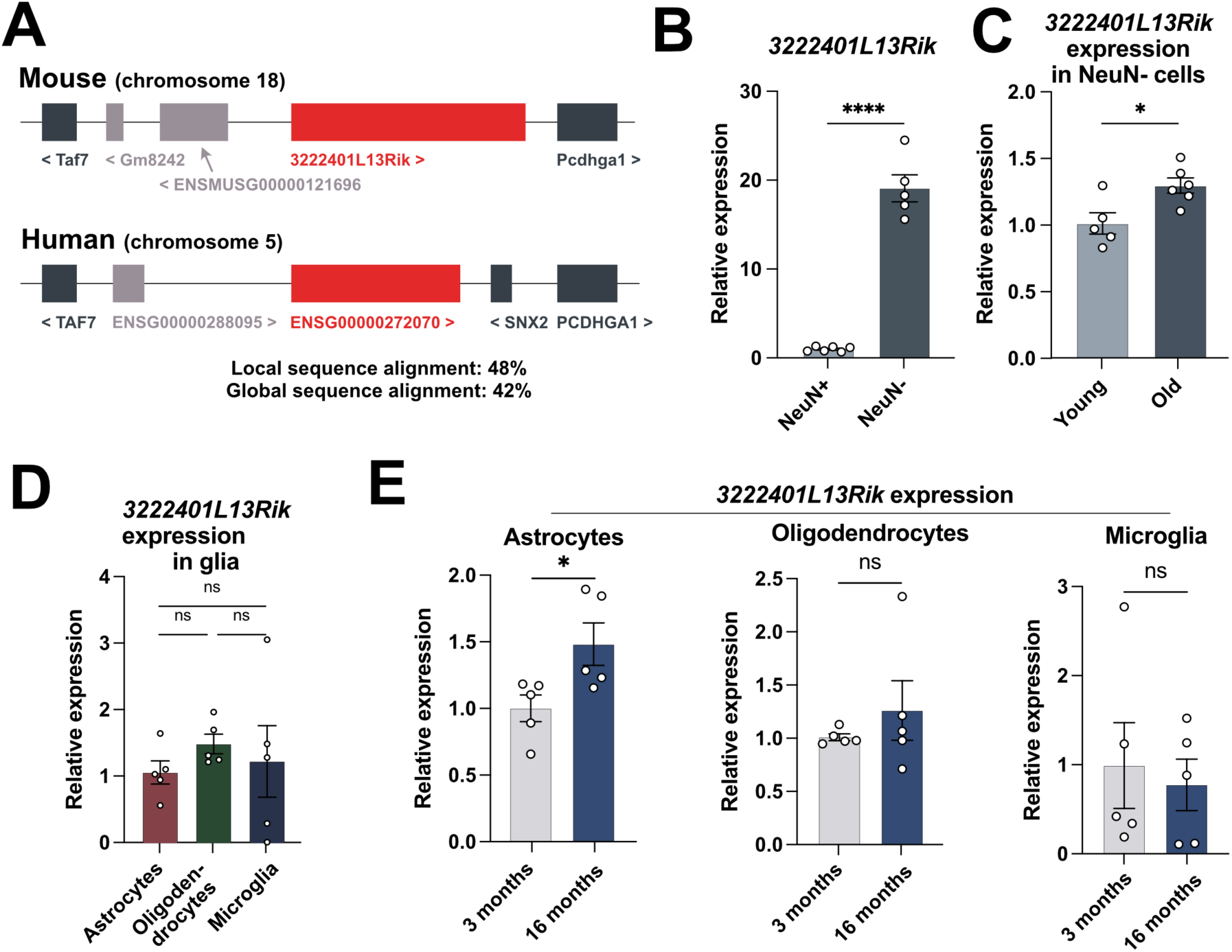
*3222401L13Rik* is a glial lncRNA that is upregulated in astrocytes during aging. **(A)** Schematic illustration of the genomic localization of *3222401L13Rik* in the mouse and *ENSG00000272070* in the human genome. **(B)** Expression of the lncRNA *3222401L13Rik* in NeuN+ and NeuN-cells isolated from the hippocampal CA1 region of 3 months old mice (unpaired t-test; ****p < 0.0001). **(C)** qPCR data showing the expression of *3222401L13Rik* in NeuN-cells from 3 and 16 months old mice (unpaired t-test; *p < 0.05). **(D)** Expression of *3222401L13Rik* in astrocytes, oligodendrocytes and microglia isolated from the brains of 3 months old mice using MACS (One-way ANOVA; ns = not significant). **(E)** Expression of *3222401L13Rik* in astrocytes, oligodendrocytes and microglia isolated from the brains of 3 and 16 months old mice using MACS technology (unpaired t-test; *p < 0.05, ns = not significant). Data are depicted as mean ± standard error.

### Loss of *3222401L13Rik* leads to expression changes of genes involved in neuronal support and inflammatory processes

lncRNAs can exert a wide range of functions, which primarily depend on their location within the cell, and the identification of their subcellular localization is therefore critical for their characterization (Bridges *et al*, 2021). Although we identified *3222401L13Rik* in Neu-nuclei, this does not exclude the possibility that it could be prominently expressed in the cytoplasms. Therefore, we investigated the localization of *3222401L13Rik* by combining RNAscope for *3222401L13Rik* with immunofluorescence staining for the astrocyte marker glial fibrillary acidic protein (Gfap), and found that *3222401L13Rik* is predominantly located in the nucleus of astrocytes in the adult mouse brain **(Fig. 3A)**. This finding was confirmed by performing nuclear and cytoplasmic fractionation of primary mouse astrocytes and measuring the abundance of *3222401L13Rik* in both compartments **(Fig. 3B)**. The enrichment of *3222401L13Rik* in the nuclear compartment hints towards a role of this lncRNA in regulating gene transcription. Therefore, we performed a knockdown (KD) of *3222401L13Rik* in primary mouse astrocytes using Gapmers directed against the lncRNA and performed total RNA sequencing to identify transcriptional targets. As a negative control (NC), a Gapmer with no known target in the genome was employed. We confirmed by qPCR that the Gapmers reduced the abundance of *3222401L13Rik* by ∼75% **(Fig. 3C)**. Differential expression analysis after total RNA sequencing revealed that 321 genes were upregulated and 754 genes were downregulated upon *3222401L13Rik* KD (|log2FC| > 0.5, p-adj. < 0.05, basemean > 50) **(Fig. 3D) (Table S3)**. The upregulated genes were linked with a number of interesting Gene Ontology (GO) terms including “Response to interferon-gamma”, “Positive regulation of nervous system development” and “Leukocyte migration” or “Extracellular matrix organization” **(Fig. 3E) (Table S4)**. The downregulated genes were associated with GO terms such as “Synapse organization”, “Regulation of membrane potential”, “Synaptic vesicle cycle” and “Locomotory behavior” **(Fig. 3E) (Table S4)**. To confirm the RNA sequencing data, we performed a qPCR for several of the upregulated genes that are part of the signaling pathway “Response to interferon gamma” and downregulated genes linked to synaptic function **(Fig. 3F)**.

**Figure 3:**
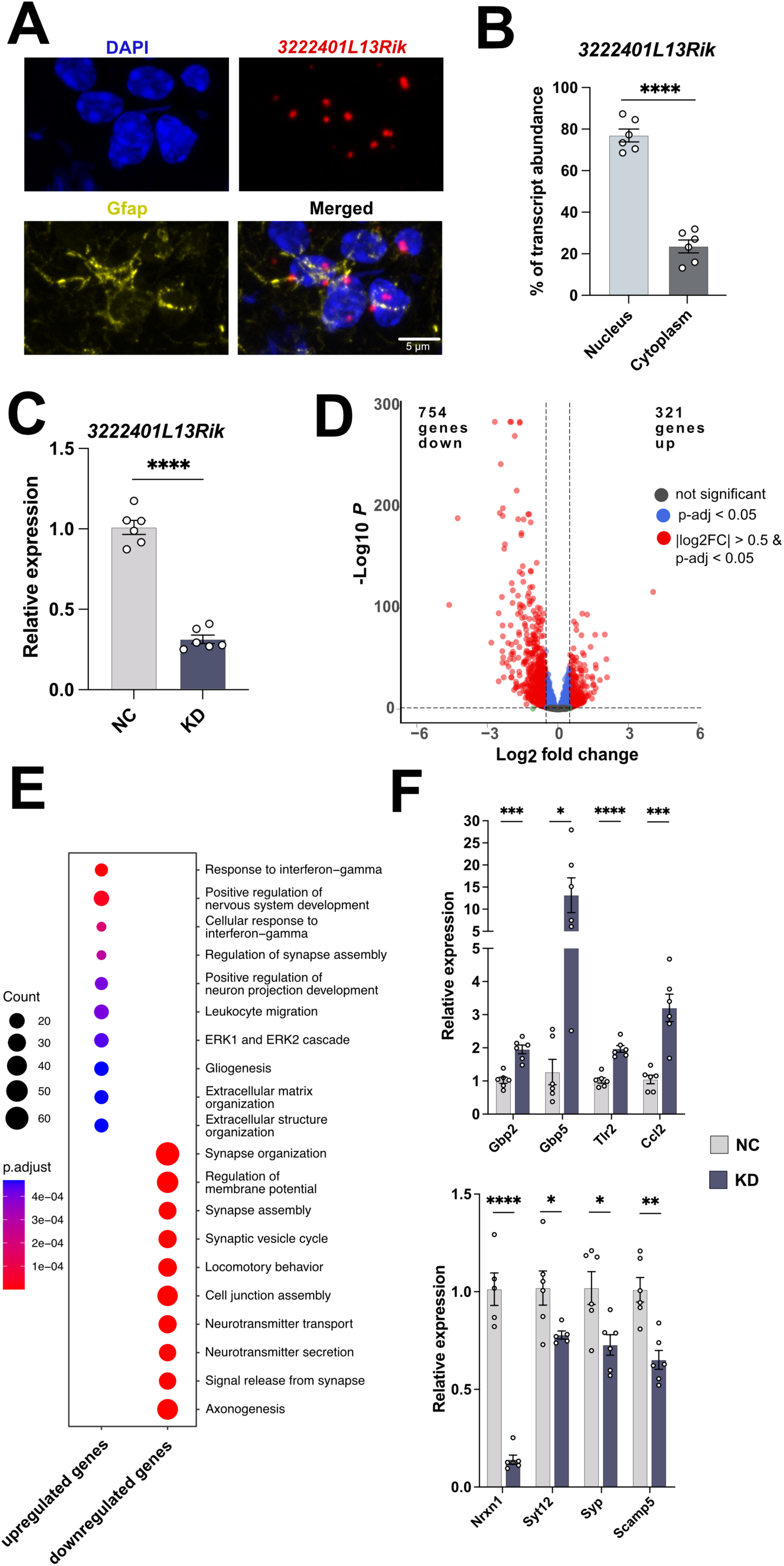
*3222401L13Rik* controls the expression of genes linked to innate immune response and synaptic support functions. **(A)** Representative image showing the nuclear localization of *3222401L13Rik* (RNAscope) in astrocytes (immunofluorescence for Gfap) in the adult mouse brain. Nuclei are stained using DAPI. **(B)** Bar chart showing the results of a qCPR that analyze the expression of *3222401L13Rik* in nuclear and cytoplasmic fractions isolated from primary astrocytes (unpaired t-test; ****p < 0.0001). **(C)** Bar charts showing qPCR results to measure the expression levels of *3222401L13Rik* after treatment with NC or KD ASOs. **(D)** Volcano Plot showing the up- and downregulated genes 48 hours after the KD of *3222401L13Rik* in primary astrocytes. **(E)** Gene Ontology analysis of the genes shown in (D). Analysis was done using clusterProfiler (v4.6.0) (Yu *et al*, 2012). Two-sided hypergeometric test was used to calculate the importance of each term and the Benjamini-Hochberg procedure was applied for the P value correction). **(F)** Expression levels of selected genes that were deregulated after the KD of *3222401L13Rik*. Upper panel: upregulated genes. Lower panel: downregulated genes (unpaired t-test; *p < 0.05, **p < 0.01, ***p < 0.001, ****p < 0.0001). Data are depicted as mean ± standard error. NC: negative control, KD: knockdown of *3222401L13Rik*.

### *3222401L13Rik* is important for glutamate uptake, Ca^2+^ signaling and neuronal support

Although it would be interesting to study further the up-regulated genes linked to inflammatory processes, the majority of deregulated genes was decreased after the KD of *3222401L13Rik*, suggesting that this lncRNA can act as an activator of gene transcription under physiological conditions. The pathways linked to these downregulated genes indicate a potential function of *3222401L13Rik* in mediating neuronal support. Therefore we decided to study in the context of tis work the role of *3222401L13Rik* in gene activation and the regulation of neuronal support. In this regard, an important role of astrocytes is the removal of glutamate from the extracellular space, which is mainly mediated by glutamate transporter 1 (Glt-1) and glutamate aspartate transporter (Glast). We found that the levels of both transporters were reduced on the RNA **(Fig. 4A)** and on the protein level **(Fig. 4B)** following the KD of *3222401L13Rik*. In line with this, also the uptake of glutamate from the extracellular space was impaired in primary astrocytes in the absence of *3222401L13Rik* **(Fig. 4C)**. Another important feature of astrocytes is the ability to generate transient increases of intracellular calcium (Ca^2+^) levels in response to neuronal activity, which can induce the release of various gliotransmitters, e.g. glutamate, ATP and GABA (Bazargani & Attwell, 2016). These gliotransmitters can in turn have a wide range of effects on neurons, such as the synchronization of action potential firing (Fellin *et al*, 2004) or the regulation of synaptic vesicle release (Perea, 2009). Pathological alterations of astrocytic Ca^2+^ dynamics have been described in different contexts, e.g. in aging and Alzheimer’s disease (Lin *et al*, 2007) (Kuchibhotla *et al*, 2009). Therefore, we measured Ca^2+^ levels in response to adenosine triphosphate (ATP) stimulation and found that the increase in intracellular Ca^2+^ was reduced after the KD of *3222401L13Rik* compared to astrocytes treated with NC Gapmers **(Fig. 4D)**.

**Figure 4:**
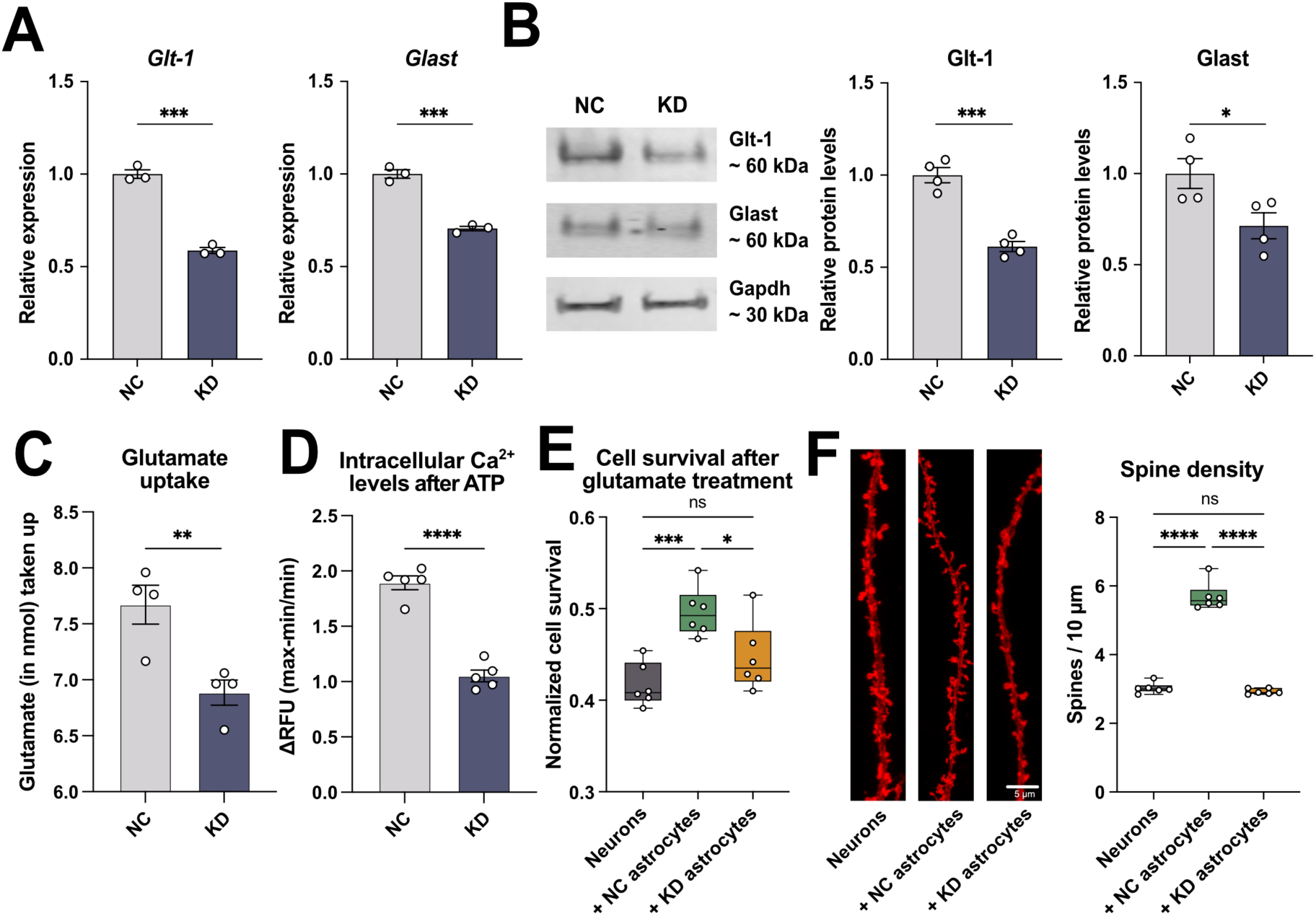
The KD of *3222401L13Rik* affects glutamate uptake, Ca^2+^ signaling and the support of neuronal function. **(A)** qPCR showing the expression levels of the glutamate transporters *Glt-1* and *Glast* after the KD of *3222401L13Rik* in primary astrocytes (unpaired t-test; ***p < 0.001). **(B)** Left panel: Representative immunoblot images of Glt-1 and Glast following the KD of *3222401L13Rik*. in primary astrocytes Right panel: Quantification of the left panel (unpaired t-test; *p < 0.05, ***p < 0.001). **(C)** Glutamate uptake of primary astrocytes after the KD of *3222401L13Rik* (unpaired t-test; **p < 0.01). **(D)** Increase in intracellular Ca^2+^ levels in response to ATP treatment after the KD of *3222401L13Rik* (unpaired t-test; ****p < 0.0001). **(E)** Survival of neurons after treatment with 100 µM glutamate cultured alone or co-cultured with NC or KD astrocytes (One-way ANOVA; *p < 0.05, ***p < 0.001, ns = not significant). **(F)** Left panel: Representative images of dendrite and spine labeling of neurons cultured alone or co-cultured with NC or KD astrocytes. Right panel: Quantification of spines shown in the left panel (One-way ANOVA; ****p < 0.0001, ns = not significant). Data are depicted as mean ± standard error. NC: negative control, KD: knockdown.

Since our previous results suggested that the lncRNA *3222401L13Rik* is important for maintaining neuronal support functions of astrocytes, we next asked whether the loss of *3222401L13Rik* in astrocytes affects neurons. Therefore, we cultured primary neurons alone or together with astrocytes treated with either control (+ NC astrocytes) or *3222401L13Rik* KD Gapmers (+ KD astrocytes). Then, we treated all groups with 100 µM glutamate and measured neuronal cell viability 3 hours later. We found that the co-culture with NC astrocytes increased neuronal viability compared to the neuronal mono-culture, most likely due to the uptake of extracellular glutamate **(Fig. 4E)**. In contrast, the co-culture with KD astrocytes did not have these beneficial effects on cell survival **(Fig. 4E)**. Moreover, we assessed the effect of NC or KD astrocyte co-cultures on neuronal spine density. We found that only NC, but not KD, astrocytes increased the number of dendritic spines **(Fig. 4F)**, suggesting that the lncRNA *3222401L13Rik* indeed plays an important role in maintaining neuronal support. This interpretation is further supported by electrophysiological measurements using Multielectrode array (MEA) assays **(Fig. S1)**.

### The regulation of synaptic support genes is conserved in human iPSC-derived astrocytes

As *3222401L13Rik* has a human homolog, we next asked whether its function is conserved between species. We performed a KD of *ENSG00000272070* in human iPSC-derived astrocytes **(Fig. 5A)** and measured via qPCR the levels of interferon response genes, which had been found to be upregulated in mouse astrocytes. However, *GBP2* and *TLR2* were significantly downregulated after the KD of *ENSG00000272070*, whereas the levels of *GBP5* and *CCL2* remained unchanged **(Fig. 5B)**. In contrast, we could recapitulate the findings from the mouse astrocytes regarding the downregulation of several genes linked to synaptic function **(Fig. 5C)** and the two glutamate transporters *GLT-1* and *GLAST* in human iPSC-derived astrocytes **(Fig. 5D)**. Furthermore, also the uptake of extracellular glutamate and the increase in intracellular Ca^2+^ after ATP stimulation were impaired after the KD of *ENSG00000272070* **(Fig. 5E, F)**. These findings suggest a functional conservation of *3222401L13Rik* and *ENSG00000272070* regarding its role in maintaining the expression of genes important for neuronal support, whereas the upregulation of interferon response genes was specific for mouse astrocytes, also supporting our decision to focus our functional analysis on the role of *3222401L13Rik* in neuronal support.

**Figure 5:**
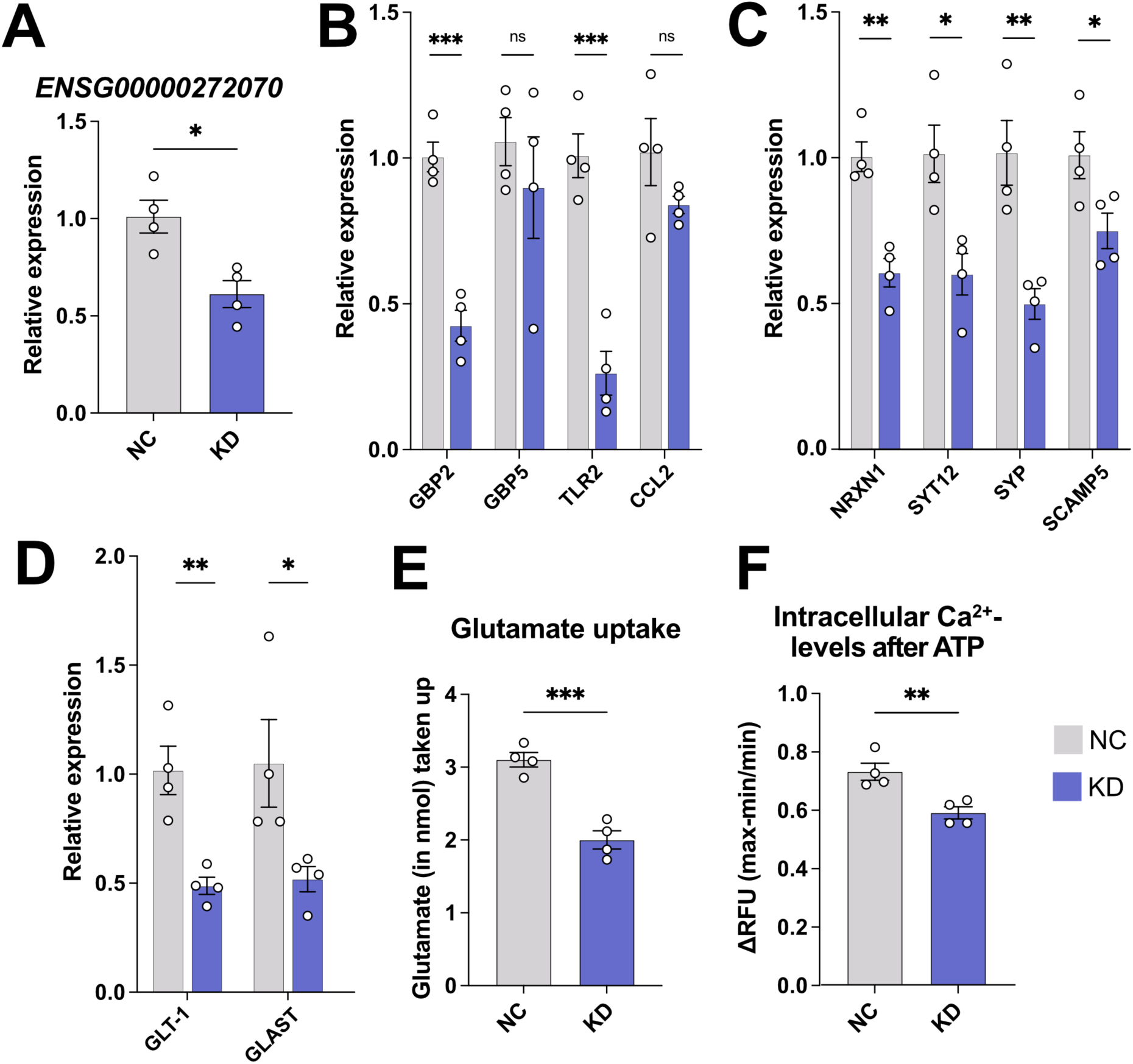
The synaptic support functions of *3222401L13Rik* are conserved in human astrocytes. **(A)** KD of *ENSG00000272070* in human iPSC-derived astrocytes (unpaired t-test; *p < 0.05). **(B)** Expression levels of interferon response genes after the KD of *ENSG00000272070* in human iPSC-derived astrocytes (unpaired t-test; ***p < 0.001, ns = not significant). **(C)** Expression levels of genes associated with synaptic support after the KD of *ENSG00000272070* in human iPSC-derived astrocytes (unpaired t-test; *p < 0.05, **p < 0.01). **(D)** qPCR showing the expression of the glutamate transporters *GLT-1* and *GLAST* after the KD of *ENSG00000272070* in human iPSC-derived astrocytes (unpaired t-test; *p < 0.05, **p < 0.01). **(E)** Glutamate uptake after the KD of *ENSG00000272070* (unpaired t-test; ***p < 0.001). **(F)** Increase in intracellular Ca^2+^ levels in response to ATP stimulation after the KD of *ENSG00000272070* (unpaired t-test; **p < 0.01). Data are depicted as mean ± standard error. NC: negative control, KD: knockdown.

### *3222401L13Rik* interacts with Npas3 to control gene expression

Among the most significantly downregulated genes identified from the RNA sequencing data was the transcription factor Neuronal PAS Domain Protein 3 (Npas3) **(Table S3)**. We confirmed this finding by qPCR in primary mouse and human iPSC-derived astrocytes **(Fig. 6A)**. Npas3 is known as an important transcription factor in astrocytes that mediates the expression of genes involved in brain development and synapse function (Li *et al*, 2022). This study also demonstrated that an astrocyte-specific knockout of Npas3, similar to our data, causes deficits in synaptic density and complexity, along with behavioral abnormalities in mice. Additionally, Npas3 target genes were found to be significantly enriched in genes associated with schizophrenia, autism, and intellectual disability (Michaelson *et al*, 2017). Based on these findings, we hypothesized that the effects observed after the knockdown (KD) of 3222401L13Rik might, at least in part, be mediated by the deregulation of Npas3.

**Figure 6:**
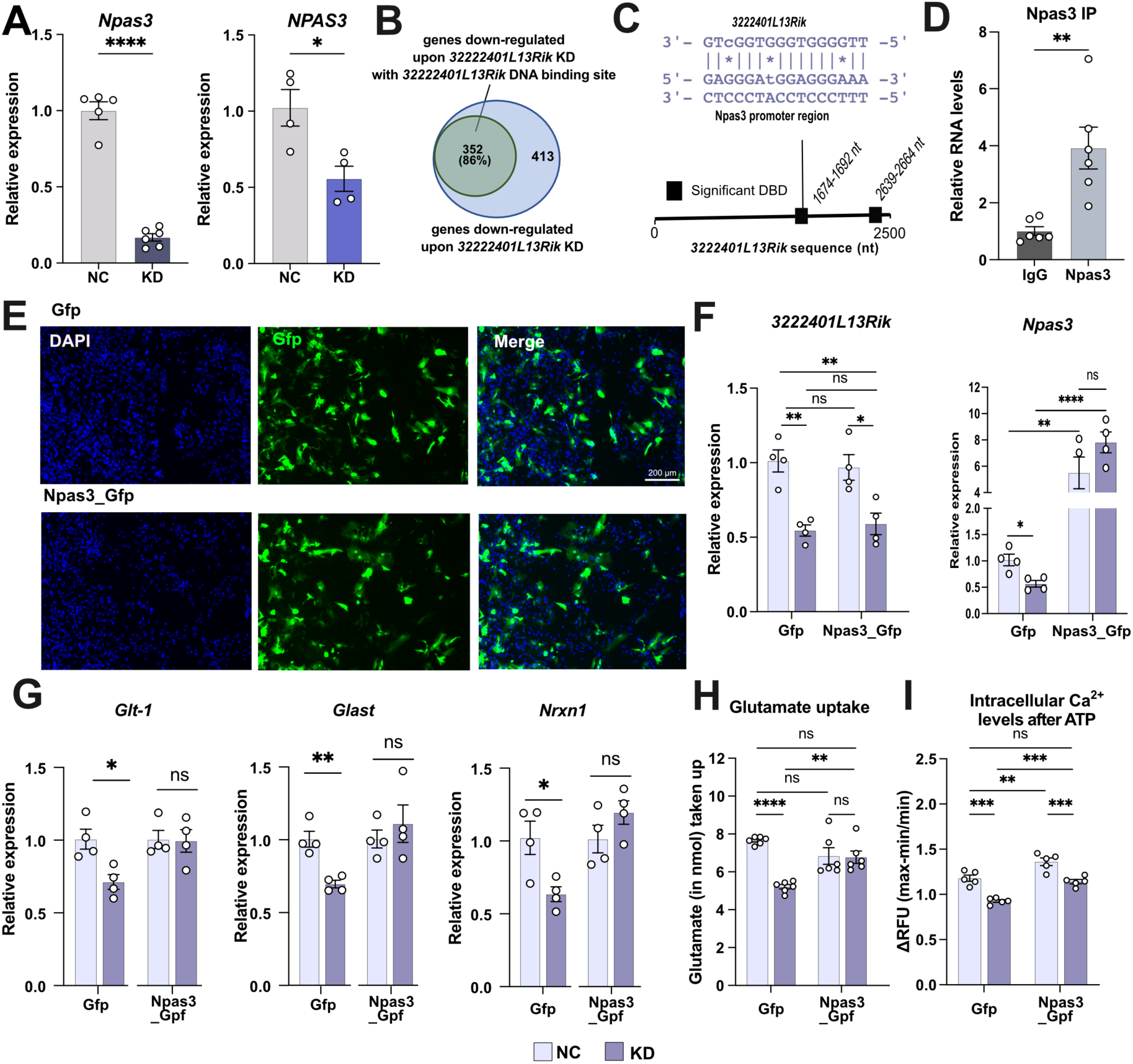
The overexpression of the interaction partner Npas3 can rescue molecular and functional changes induced by the loss of *3222401L13Rik*. **(A)** qPCR showing the expression levels of *Npas3/NPAS3* in mouse (left panel) and human iPSC-derived (right panel) astrocytes after the KD of *3222401L13Rik* (unpaired t-test; *p < 0.05, ****p < 0.0001). (**B)** Venn diagram showing the proportion of down-regulated genes (352 out of 765) containing a promoter region that can bind *3222401L13Rik*. The Triplex Domain Finder tool (Kuo *et al*., 2019) was used to identify promoter regions of the down-regulated genes that have a binding motif for *3222401L13Rik*. (**C)** Scheme depicting the significant DNA binding domains (DBD) in the sequence of *3222401L13Rik* determined using the Triplex Domain Finder tool and the sequence motifs where *3222401L13Rik* binds to the promoter of Npas3. **(D)** RNA immunoprecipitation for Npas3 followed by qPCR for *3222401L13Rik* in mouse primary astrocytes (unpaired t-test; **p < 0.01). **(E)** Representative immunofluorescence images showing the transfection of primary astrocytes with Gfp- or Gfp-Npas3-overexpression plasmids. **(F)** Expression levels of *3222401L13Rik* and Npas3 after the simultaneous KD of *3222401L13Rik* and overexpression of Npas3 in primary astrocytes (One-way ANOVA; *p < 0.05, **p < 0.01, ****p < 0.0001, ns = not significant). **(G)** Expression levels of *Glt-1*, *Glast* and *Nrxn1* after the simultaneous KD of *3222401L13Rik* and overexpression of Npas3 in primary astrocytes (One-way ANOVA; *p < 0.05, **p < 0.01, ns = not significant). **(H)** Glutamate uptake of primary astrocytes after the simultaneous KD of *3222401L13Rik* and overexpression of Npas3 (One-way ANOVA; **p < 0.01, ****p < 0.0001, ns = not significant). **(I)** Increase in intracellular Ca^2+^ levels in response to ATP stimulation after the simultaneous KD of *3222401L13Rik* and overexpression of Npas3 (One-way ANOVA; **p < 0.01, ***p < 0.001, ns = not significant). Data are depicted as mean ± standard error. NC: negative control, KD: knockdown.

lncRNAs are known to bind to DNA and form triple helices, acting as scaffolds to facilitate interactions between proteins, RNA, and DNA (Kuo *et al*, 2019). We employed the Triplex Domain Finder tool (Kuo, 2019) to assess how many of the downregulated genes can potentially bind *3222401L13Rik* via their promoter region. This analysis revealed that 46% (352 out of 765 transcripts) exhibit *3222401L13Rik* binding regions **(Fig 6B)** that are localized towards the 3’ end of *3222401L13Rik* **(Fig. 6C)**, including the potential binding site for the *Npas3* promot**er (Fig 6C),** further supporting our biochemical data suggesting that *3222401L13Rik* can directly bind to the promoter of *Npas3*. It is known that *Npas3* expression is regulated by autoregulation, where the Npas3 protein binds to its own gene promoter to drive its expression (Michaelson *et al*., 2017). Therefore, we hypothesized that *3222401L13Rik* might interact with both the *Npas3* promoter and Npas3 itself to regulate its expression. Indeed, we observed a direct interaction between *3222401L13Rik* and Npas3, as demonstrated by RNA immunoprecipitation followed by qPCR **(Fig. 6D)**.

These data support a model in which *3222401L13Rik* positively regulates *Npas3* expression, likely by promoting positive autoregulation. Loss of *3222401L13Rik* reduces Npas3 levels. To test whether overexpression of *Npas3* could rescue the functional impairments observed after *3222401L13Rik* knockdown, we electroporated primary mouse astrocytes with plasmids containing *Npas3* and *Gfp* (Npas3_GFP) under the control of the astrocyte-specific GFAP promoter - to monitor successful transfection - or only Gfp as a control (Gfp_control) **(Fig. 6E)**. We then combined this with the transfection of Gapmers to knock down *3222401L13Rik* or control Gapmers. While *3222401L13Rik* knockdown reduced the expression of Npas3 in astrocytes transfected with GFP **(Fig. 6F)**, we observed significant overexpression of Npas3 in astrocytes electroporated with the Npas3_GFP construct **(Fig. 6F)**. Importantly, knockdown of *3222401L13Rik* did not affect *Npas3* expression in this overexpression setting **(Fig. 6F)**, allowing us to test if Npas3 overexpression could rescue the phenotypes associated with loss of *3222401L13Rik* function.

Consistent with previous data (see Figs. 3F & 4A), knockdown of *3222401L13Rik* reduced the levels of *Glt-1, Glast*, and *Nrxn1* **(Fig. 6G)** in astrocytes electroporated with the GFP plasmid. However, overexpression of Npas3 reinstated physiological expression levels of all three genes despite the knockdown of *3222401L13Rik* **(Fig. 6G)**. Similar observations were made for glutamate uptake and intracellular calcium levels. Consistent with previous data (see Fig. 4C), glutamate uptake was impaired upon *3222401L13Rik* knockdown in astrocytes transfected with GFP **(Fig. 6G)**, while *Npas3* overexpression could rescue this phenotype **(Fig. 6H)**. Finally, we measured intracellular Ca2+ levels following ATP stimulation. Similar findings were observed for intracellular Ca2+ levels upon ATP stimulation **(Fig. 6I)**.

In summary, these data suggest that *3222401L13Rik* positively regulates and fine-tunes the expression of Npas3, and that the phenotypes observed upon 3222401L13Rik knockdown are, at least in part, mediated by loss of Npas3 function.

## Discussion

In this study, we identify *3222401L13Rik* as a glial-enriched lncRNA with elevated expression levels in astrocytes of aged mice. To our knowledge, the function of this lncRNA has not been studied in any cellular system before. However, *3222401L13Rik* was found to be upregulated in two distinct RNA-seq datasets related to brain aging. Specifically, upregulation of *3222401L13Rik* was observed in ApoD knockout mice, which are characterized by premature brain aging, when bulk cortical tissue was analyzed via RNA-seq (Sanchez *et al*, 2015). Similarly, upregulation of *3222401L13Rik* was detected in the hippocampi of young and old mice, as characterized by both bulk and single-cell RNA-seq (Pérez *et al*, 2023). Although neither study specifically highlights *3222401L13Rik*, these findings support our observation and suggest a role for *3222401L13Rik* in brain aging.

We found that *3222401L13Rik* is highly expressed in glial cells compared to neurons, which aligns with previous research showing that lncRNAs often exhibit tissue-specific expression patterns (Derrien *et al*., 2012). The age-associated upregulation of *3222401L13Rik* was specific to astrocytes. While we cannot yet explain the underlying molecular mechanisms, it is known that stimuli such as environmental enrichment, aging, and neurodegenerative diseases lead to cell type-specific changes in gene expression in the brain (Methi *et al*, 2024) (Mathys *et al*, 2018). Moreover, astrocytes play a crucial role in brain aging, and recent studies suggest that transcriptional changes in astrocytes are a central mechanism for promoting resilience against cognitive decline in Alzheimer’s disease (Mathys *et al*, 2024). Therefore, we decided to study the function of *3222401L13Rik* in astrocytes. It is likely that *3222401L13Rik* also regulates important cellular processes in other glial cells, such as microglia and oligodendrocytes, which we plan to address in future research.

We found that *3222401L13Rik* is primarily expressed in the nucleus, which aligns with the known role of lncRNAs in gene expression control (Mattick *et al*., 2023). Consistent with this, we observed that knockdown of *3222401L13Rik* in primary mouse astrocytes led to significant changes in gene expression, including an upregulation of inflammatory processes and a downregulation of genes associated with neuronal support functions. This finding is consistent with previous studies showing that lncRNAs can have profound effects on astrocytic gene expression control (Schröder *et al*., 2024) (Irwin *et al*., 2023).

In human iPSC-derived astrocytes, knockdown of the homologous lncRNA *ENSG00000272070* also decreased the expression of genes linked to neuronal support functions. However, no increased expression of genes linked to inflammatory processes was observed. It is important to note that this data is based on qPCR analysis of selected transcripts. Nonetheless, recent studies comparing the transcriptomic signatures of astrocyte reactivity in rodent and human models have identified both commonalities and species-specific gene signatures (Li *et al*, 2021) (Brase *et al*, 2023). Thus, the differences observed in our study may reflect partially altered regulation of astrocyte reactivity between mouse and human astrocytes.

Interestingly, previous studies using postmortem human brain tissue from AD patients have suggested a modest increase in proinflammatory genes in astrocytes and, more strikingly, a significant decrease in the expression of homeostatic genes involved in synapse regulation. Based on these findings, we further characterized the role of *3222401L13Rik* and *ENSG00000272070* in neuronal support functions. However, we acknowledge the importance of elucidating the role of these lncRNAs in inflammatory processes in future research.

Our findings demonstrate that *3222401L13Rik*, as well as its human homolog, play crucial roles in maintaining the expression of several genes critical for neuronal and synaptic support within astrocytes. These include the glutamate transporters Glt1 and Glast, as well as genes controlling synaptic plasticity such as Nrxn1. Consistent with this, knockdown of *3222401L13Rik* impaired glutamate uptake and regulation of intracellular Ca²⁺ levels in astrocytes. Additionally, astrocytes lacking *3222401L13Rik* were unable to efficiently support neurons, as evidenced by altered neuronal network plasticity and decreased neuronal spine density. Although specific data for *3222401L13Rik* or ENSG00000272070 are currently lacking, previous studies have shown that lncRNAs can regulate key cellular functions in astrocytes, including glutamate metabolism, calcium signaling, and neuronal network homeostasis (Schröder *et al*., 2024) (Irwin *et al*., 2023) (Zheng *et al*, 2023) (Nassar *et al*, 2022).

One mechanism by which lncRNAs control gene expression is through interactions with DNA, particularly in gene-promoter regions, and with proteins, thereby orchestrating the function of transcriptional regulators (Mattick *et al*., 2023). Bioinformatic analysis revealed that half of the transcripts downregulated upon *3222401L13Rik* knockdown contain binding regions for *3222401L13Rik* in their promoter regions, suggesting that these may be direct targets. Future work will be required to confirm these DNA-lncRNA interactions experimentally. Among these potential targets is Npas3, a transcription factor associated with neuropsychiatric disorders characterized by impaired neuronal plasticity, such as schizophrenia (Kamnasaran *et al*, 2003) (Lavedan *et al*, 2009) (Macintyre *et al*, 2010) (Brunskill *et al*, 2010) (Erbel-Sieler *et al*, 2004). Our study shows that *3222401L13Rik* regulates *Npas3*, which is noteworthy since *Npas3* is highly expressed in astrocytes, and its deficiency induces synaptic deficits in neurons (Li *et al*., 2022). Moreover, the affected pathways and gene networks identified in the study closely resemble the downregulated genes and pathways observed after knockdown of *3222401L13Rik*, including terms like “synapse organization,” “neurotransmitter transport,” and “synapse assembly.” Accordingly, we demonstrate that overexpression of *Npas3* can rescue both molecular and functional deficits resulting from knockdown of *3222401L13Rik*. These data strongly suggest that *3222401L13Rik* orchestrates the regulation of astroglia function - at least in part - via Npas3.

Taking into account that *3222401L13Rik* expression increases in the aging brain, and that its knockdown impairs astrocytes’ ability to perform critical neuronal support functions, it can be speculated that altered *3222401L13Rik* expression may represent an adaptive mechanism aimed at mitigating ongoing network imbalances and functional alterations associated with aging. However, further studies are needed to determine whether modulating *3222401L13Rik* and *ENSG00000272070* levels can indeed enhance astrocytic function and neuronal support during aging.

Our study has several limitations that should be acknowledged. First, increasing evidence reveals distinct astrocyte subtypes within and between brain regions (Batiuk *et al*, 2020). Since our study focused on the hippocampus, it will be important to investigate the role of *3222401L13Rik* and *ENSG00000272070* in other brain regions. Additionally, analyzing the expression of these lncRNAs in postmortem human brains from both young and old individuals, as well as those with cognitive diseases, will be crucial. Ideally, this should be complemented by similar analyses in animal models, allowing us to determine if *3222401L13Rik* expression is a compensatory mechanism aimed at ameliorating the detrimental effects of brain aging. This is particularly interesting given recent data suggesting a key role of astrocytes in mediating resilience to cognitive decline (Mathys *et al*., 2024). It would also be valuable to study the functional role of *3222401L13Rik* in an in vivo system. Knockdown of *3222401L13Rik* in wild-type mice and its overexpression in disease models could provide important insights critical for the preclinical development of RNA-based cognitive enhancers. However, preclinical research involving mice is becoming increasingly difficult in the European Union, particularly in Germany, and we are unable to perform such experiments within a reasonable time frame. Finally, our dataset suggests several other lncRNAs that are deregulated in the aging mouse brain and have human homologues. In addition to *Neat1*, which is well-studied for its role in brain function and disease (Irwin *et al*., 2023) (Butler *et al*, 2019) (Zhao *et al*, 2020) (Yamada *et al*, 2022) (Yadav *et al*, 2024), we identified *A330023F24Rik*, *Lncpint*, *Gm12689*, *Mir124-2hg*, *B130024G19Rik*, *C430049B03Rik*, and *Gm45847* as being deregulated in the aging mouse hippocampus and having human homologues. *A330023F24Rik* is deregulated in Duchenne muscular dystrophy (Xu & Zheng, 2022) and in response to optic nerve injury (Sun *et al*, 2024). *Lncpint*, a neuron-enriched lncRNA, is increased in various neurodegenerative diseases, including Alzheimer’s disease (AD), Parkinson’s disease, and Huntington’s disease and functional studies suggest a protective role, since loss of Lncpint exacerbates pathology (Simchovitz *et al*, 2020), which is similar to our data on *3222401L13Rik. Mir124-2hg* is also a neuron-enriched lncRNA with a reported role in synaptic vesicle recycling (Keihani *et al*, 2019) that warrants further analysis in neurons. The human homologue of *B130024G19Rik, NR2F2-AS1,* has been studied in various cancers, as has *C430049B03Rik* and its human homologue *MIR503HG*.

In summary, our study identifies the novel lncRNA *3222401L13Rik* as crucial in regulating astrocytic gene expression through its interaction with the transcription factor Npas3. The observed upregulation of *3222401L13Rik* suggests that it acts as an adaptive mechanism in astrocytes to support neuronal function during aging. Given that age-associated memory decline and the pathogenesis of neurodegenerative diseases are characterized by distinct molecular and cellular phases (De Strooper & Karran, 2016), strategies aimed at enhancing *3222401L13Rik* function could offer promising avenues for developing targeted therapies for cognitive diseases.

## Material and methods

### Animals

All animal experiments were approved by the local Animal Welfare Office of Goettingen University and the Lower Saxony State Office for Consumer Protection and food safety. Three and 16 months old male C57/BL6 and pregnant CD-1 mice were obtained from Janvier Labs. Animals were housed in standard cages with a 12-hour dark and light cycle. Water and food were provided ad libitum.

### Sorting of neuronal and non-neuronal nuclei from mouse brain

Young and old C57/BL6 mice were sacrificed using pentobarbital or CO_2_. Brains were quickly removed and the CA1 region of the hippocampus was dissected using a dissection microscope. Brain tissue was flash-frozen and kept at -80°C until further use. Nuclei isolation, NeuN-staining and FANS-sorting was performed as previously described (Sakib *et al*, 2021) (Michurina *et al*, 2022). After sorting, Trizol LS (ThermoFisher) was added, and RNA was isolated as described below.

### Primary astrocyte culture

For the generation of primary astrocyte cultures, a protocol was used that allows the cultivation of astrocytes that are less reactive compared to cultures permanently containing fetal bovine serum (FBS) (Wolfes & Dean, 2018). Briefly, postnatal day 0 to 2 pups were sacrificed by decapitation, the brains quickly taken out and the meninges removed. Cortices and hippocampi were dissociated using 0.05% Trypsin-Ethylenediaminetetraacetic acid (EDTA) (Gibco), and the resulting cells were seeded into T75 flasks coated with 0.5 mg/mL Poly-D-lysine (PDL) (MerckMillipore) and kept in DMEM containing 1% penicillin-streptomycin and 10% FBS (all Gibco) in a humidified incubator with 5% CO_2_ at 37°C. After 7-8 days, the flasks were shaken at 160 rpm for 6 hours to remove non-astrocytic cells. Astrocytes were removed from the flasks using 0.25% Trypsin-EDTA (Gibco), centrifuged, counted and seeded at a density of 15,000 cells/cm^2^ on pre-coated cell culture dishes. Cells were kept in Neurobasal Plus Medium containing 2% B27 Plus supplement, 1x GlutaMax and 1% penicillin-streptomycin (NB+) (all Gibco) with 5 ng/mL heparin-binding EGF-like growth factor (HB-EGF) (Sigma-Aldrich) until experiments were performed.

### Primary neuron culture

Pregnant CD-1 mice were sacrificed using pentobarbital on embryonic day 17. The embryos were taken out and their brains removed. After dissection of the meninges, cortices and hippocampi were dissociated using the Papain Dissociation System (Worthington) as described by the manufacturer’s instructions. Finally, cells were resuspended in NB+, counted and seeded at a density of 120,000 cells/cm^2^ for glutamate treatment experiments or 60,000 cells/cm^2^ for spine analysis on cell culture dishes pre-coated with 0.5 mg/mL PDL or 0.1% polyethylenimine (PEI).

On day in vitro (DIV) 14, astrocytes cultured in co-culture inserts (Greiner) were added to neurons and experiments were performed on DIV17.

### Human iPSC-derived astrocytes

Human induced pluripotent stem cell (iPSC)-derived astrocytes were obtained from Ncardia. Cells were shipped and stored in liquid nitrogen until use. Cells were thawed and cultured according to the manufacturer’s instructions. Transfections and experiments were performed at least one week after thawing.

### Antisense LNA Gapmers

Custom Antisense LNA Gapmers directed against *3222401L13Rik* and *ENSG00000272070* as well as negative controls were designed by and obtained from Qiagen having the following sequences:

NC: GCTCCCTTCAATCCAA

*3222401L13Rik*: AGCTTGGTCATTTGAT

*ENSG00000272070*: GGACTTCTTCCTCTGT

Primary and iPSC-derived astrocytes were transfected with 50 nM Gapmers using Lipofectamine RNAiMax (ThermoFisher) according to the manufacturer’s instructions. Primary astrocytes were transfected on DIV12, and functional experiments were performed 48 hours later. iPSC-derived astrocytes were transfected at least one week after thawing and collected after 48 hours as well.

### Npas3 overexpression

Expression plasmids encoding the open reading frame for Npas3-Gfp or Gfp only under the control of the glial fibrillary acidic protein (Gfap)-promoter were designed by and purchased from VectorBuilder. Primary astrocytes were transfected on DIV7 before seeding using the Neon Nxt electroporation device (ThermoFisher). 100,000 cells were electroporated using 0.25 µg plasmid and 25 nM Antisense LNA Gapmers (1300 V, 20 ms, 2 pulses) and seeded in pre-coated 24 well plates. Experiments were performed 72 hours after transfection.

### Glutamate uptake

To measure the amount of glutamate taken up from the extracellular space, primary and iPSC-derived astrocytes were pre-incubated with Hank’s Buffered Salt Solution (HBSS) for 10 min at 37°C. Then, the supernatant was aspirated and 100 µM glutamate in HBSS added. After 1 hour for primary astrocytes and 3 hours for iPSC-derived astrocytes, the supernatant was collected, and the remaining glutamate was measured using the Glutamate-Glo™ Assay (Promega) according to the manufacturer’s instructions. Luminescence was recorded with a FLUOstar® Omega plate reader (BMG).

### Measurement of intracellular Ca^2+^ levels

Primary mouse and human iPSC-derived astrocytes were incubated with Fluo-4 calcium assay reagent (Fluo-4 Direct™ Calcium-Assay-Kit, ThermoFisher) for 30 min at 37°C before recording baseline fluorescence levels (RFU_Min_) using a plate reader. Then, 100 µM ATP were added and the plate was measured immediately again (RFU_Max_). The difference in relative fluorescence units (ΔRFU) was calculated using the following formula: ΔRFU=(RFU_Max_-RFU_Min_)/RFU_Min_. As a blank, wells containing no cells were used.

### Collection of astrocyte-conditioned medium (ACM)

For the treatment of neurons with ACM, medium from NC or KD astrocytes was collected 48h after treatment, filtered using a 0.22 µm filter and stored at -80°C until use. On DIV14, primary neurons were treated with ACM that was diluted 1:1 with fresh NB+.

### Glutamate treatment of primary neurons

Primary neurons cultured either alone or together with NC or KD astrocytes or with ACM from NC or KD astrocytes were treated with 100 µM glutamate for 15 minutes at 37°C. Then, the medium was removed and fresh NB+ added. Three hours later, cell viability was assessed using PrestoBlue (ThermoFisher).

### Multielectrode array

Multielectrode array (MEA) recordings were performed in the Maestro system (Axion) as previously described (Castro-Hernández *et al*, 2023). ACM from NC or KD astrocytes was added on DIV14 and recordings were performed twice a day until DIV17.

### Dendrite and spine analysis

Spines from primary neurons cultured either alone or together with NC or KD astrocytes or with NC or KD ACM were analyzed as previously described (Cheng *et al*, 2014). Briefly, cells were fixed using 2% paraformaldehyde (PFA), and dendrites and spines were labeled using the dye 1,1ʹ-dioctadecyl- 3,3,3ʹ,3ʹ-tetramethylindocarbocyanine perchlorate (Dil) (ThermoFisher). Dendrite length and the number of spines were measured using ImageJ software.

### Sorting of astrocytes, oligodendrocytes and microglia using MACS

Three- and 16-months old C57/BL6 mice were sacrificed using pentobarbital or CO_2_. The brains were quickly isolated, and the meninges were removed by carefully rolling the brains over Whatman paper. Then, the tissue was dissociated using the adult brain dissociation kit (cat. no. 130-107-677, Miltenyi) according to the manufacturer’s protocol with minor modifications. Briefly, the minced tissue was incubated with the enzyme mix for 30 minutes at 37°C in a water bath and gently triturated after 5, 15 and 25 minutes. Then, the suspension was applied to 40 µm cell strainers, and the steps for red blood cell and debris removal were performed. Antibody labeling was done using the following antibodies: Anti-O4 microbeads (1:40, cat. no. 130-094-543), Anti-ACSA2 microbeads (1:10, cat. no. 130-097-678) and Anti-Cd11b microbeads (1:10, cat. no. 130-093-634). Oligodendrocytes, astrocytes and microglia were isolated using the MACS technique. The purity of the resulting populations was assessed using qPCR.

### RNAscope combined with immunofluorescence

To stain for both *3222401L13Rik* and the astrocyte marker Gfap, we combined RNAscope Fluorescent Multiplex assays (Acdbio) and immunofluorescence according to the manufacturer’s protocol for fresh frozen tissue. Briefly, C57/BL6 mice were sacrificed using pentobarbital or CO_2_, and the brains were quickly removed, embedded in optimal cutting temperature (OCT) compound, flash-frozen in liquid nitrogen and stored at -80°C until use. 18 µm tissue sections were prepared using a cryostat (Leica). Then, the tissue sections were fixed using 10 % neutral buffered formalin, dehydrated and treated with hydrogen peroxide before the overnight incubation with the primary antibody anti-Gfap (rabbit, Abcam; 1:250). On the next day, after a post-primary fixation, *3222401L13Rik* was labeled using the RNAscope® Multiplex Fluorescent Reagent Kit v2 (Acd Bio) with probes designed to target the lncRNA and TSA Plus Cyanine 5 (Akoya Biosciences; 1:750) for detection. Finally, the secondary antibody (Alexa Fluor™ 555 goat anti-rabbit secondary antibody; 1:1000, ThermoFisher) and DAPI (Sigma-Aldrich) were applied, and the slides were mounted using Prolong Gold Antifade Reagent (ThermoFisher). Confocal images were acquired within a week after staining.

### Imaging

Fluorescent images were taken with a Leica dmi8 microscope, confocal images with the same microscope fitted with a STEDycon STED/Confocal (Abberior) in confocal mode, with a 63X or 100X oil immersion objective.

### Cytoplasmic and nuclear fractionation

Primary astrocytes were harvested on DIV14 using 0.25% Trypsin-EDTA, and the resulting pellets were washed with PBS. To isolate the cytoplasmic fraction, 500 µL EZ prep lysis buffer (Sigma-Aldrich) supplemented with RNase inhibitor (Promega) was added and incubated on ice for 7 minutes. The samples were centrifuged, and the supernatant, containing the cytoplasmic fraction, was collected. The nuclear pellet was washed with EZ prep lysis buffer and resuspended in 1.5 mL phosphate buffered saline (PBS) supplemented with 0.5% bovine serum albumin (BSA) (CellSignal), RNase inhibitor and protease inhibitor (Roche). After centrifugation, the supernatant was aspirated, leaving 250 µL of liquid in the tube. TRIzol™ LS Reagent (ThermoFisher) was added to both the cytoplasmic and nuclear fractions, and RNA was extracted as described in the next section.

### RNA isolation

Cells were lysed using TRI reagent (Sigma-Aldrich), and subsequent RNA extraction was done using the RNA clean & concentrator-5 kit (Zymo Research) according to the manufacturer’s instructions. RNA concentrations were measured with Nanodrop or Qubit (both ThermoFisher). RNA quality was assessed using a Bioanalyzer (Agilent Technologies) before subjecting the samples to total RNA sequencing.

### cDNA synthesis and qPCR

Transcriptor cDNA first strand Synthesis Kit (Roche) and random hexamer primers were used to prepare cDNA from 20-800 ng RNA starting material according to the manufacturer’s instructions. Synthesized cDNA was diluted to a concentration of 1 ng/µL with nuclease-free water. The qPCR reactions were prepared using LightCycler 480 SYBR Master Mix (Roche) and run in duplicates in a LightCycler 480 (Roche). Primer sequences are listed in Table S5. Analysis was done using the 2^-DDCt method (Livak & Schmittgen, 2001).

### Library preparation and total RNA sequencing

The SMARTer Stranded Total RNA Sample Prep Kit - HI Mammalian kit (Takara Bio) with 300 ng RNA as input was used for library preparation according to the manufacturer’s instructions. 13 cycles were applied for library amplification, and the quality of the libraries was assessed using a Bioanalyzer. Then, the multiplexed libraries were sequenced in a NextSeq 2000 (Illumina) with a 50 bp single-read configuration.

### Bioinformatic analysis

Raw reads were processed and demultiplexed using bcl2fastq (v2.20.2). Quality control of raw sequencing data was done using FastQC (v0.11.5). Reads were aligned to the mouse (mm10) genome with the STAR aligner (v2.5.2b), and feature Counts (v1.5.1) was used to generate read counts. Differential gene expression was performed with DESeq2 (v1.38.3) (Love *et al*, 2014). For this, normalized read counts were used and correcting for unwanted variation detected by RUVSeq (v1.32.0) (Risso *et al*, 2014) was applied. GO term analysis was performed using clusterProfiler (v4.6.0) (Yu *et al*., 2012).

### Western Blot

Primary astrocytes were harvested and lysed with RIPA buffer (ThermoFisher) containing 1x protease inhibitor (Roche). Protein concentration was measured with Pierce BCA Protein Assay Kit (ThermoFisher), and 20 µg protein was used per well of a 4–20% Mini-PROTEAN® TGX™ Precast Protein Gel (Bio-Rad). Protein denaturation was carried out using 8 M Urea in 1x Laemmli Sample Buffer (Bio-Rad) for 60 minutes at 40°C. The gels were run at 90 V for 15 minutes and 120 V for 50 minutes. Transfer to low-fluorescence PVDF membranes was performed with the Trans-Blot Turbo Transfer System (both from Bio-Rad).

Then, membranes were blocked with 5 % BSA in PBS + 0.1 % Tween-20 (PBS-T), followed by overnight incubation with the following primary antibodies: anti-Eaat1 (rabbit, Abcam; 1:2500), anti-Eaat2 (rabbit, Abcam; 1:800) and anti-Gapdh (mouse, ThermoFisher; 1:4000) (Table S6). On the next day, the membranes were washed three times with PBS-T and incubated with the corresponding secondary antibodies (IRDye, LI-COR; 1:10,000) for 1 hour at room temperature. After three subsequent washes with PBS-T, the membranes were imaged with an Odyssey DLx (LI-COR), and the blots were analyzed with ImageJ software.

### RNA immunoprecipitation (RNA-IP)

Primary astrocytes were cultured in 15 cm dishes and harvested on DIV14. Cell pellets were resuspended in fractionation buffer (10 mM Tris-HCl, 10 mM NaCl, 3 mM MgCl2, 0.5% Nonidet P-40 (NP-40), 1mM DTT, 100 units/mL RNase inhibitor, 1x protease inhibitor) and incubated on ice for 10 minutes with occasional pipetting to lyse cells. Then, the samples were centrifuged at 1000 g, 5 min, 4°C, and the supernatant containing the cytoplasmic fraction was transferred into fresh tubes. The pellet was washed with TSE buffer (10 mM Tris, 300 mM sucrose, 1 mM EDTA, 0.1% NP-40, 1mM DTT, 100 units/mL RNase inhibitor, 1x protease inhibitor), resuspended in fresh TSE buffer, transferred to bioruptor tubes and sonicated in a Bioruptor Plus (both Diagenode) for ten cycles (30s on, 30s off). Afterwards, the samples were incubated on ice for 20 minutes with occasional vortexing and centrifuged at 14500 g for ten minutes. The supernatant was transferred to a fresh tube, and the protein concentration of both fractions was determined as described above. Then, all samples were flash-frozen and stored at -80°C until further use.

For the RNA-IP, 500 µg protein from the nuclear lysate per sample were pre-cleared using 25 µL Pierce™ Protein A/G magnetic beads (ThermoFisher) for 1 hour at 4°C. Simultaneously, 1.5 µg Npas3 antibody (rabbit, ThermoFisher) or 1.5 µg IgG isotype control (rabbit, ThermoFisher) per sample were incubated with 50 µL Protein A/G magnetic beads for two hours at RT, followed by a wash with RNA-IP buffer (50 mM Tris-HCl, 100 mM NaCl, 32 mM NaF, 0.5% NP-40). When pre-clearing of the samples was done, samples were put on a magnetic rack to separate the beads from the lysate, 10 % of each sample was collected to serve as an input control and the remaining liquid was added to the beads. The samples were incubated overnight at 4°C with mixing. On the next day, the beads were washed five times with RNA-IP buffer and then resuspended in a proteinase K buffer containing proteinase K (Qiagen), followed by an incubation at 37°C for one hour. The beads were again separated using a magnetic rack and the supernatant was transferred to fresh tubes. RNA was extracted using the RNA clean & concentrator-5 kit (Zymo Research) according to the manufacturer’s instructions.

### Statistical analysis

For statistical analysis, GraphPad Prism version 9 was used. All graphs are shown as mean ± standard error unless stated otherwise. A two-tailed unpaired t-test or a one-way ANOVA with Tukey’s post hoc test were applied for data analysis. Enriched gene ontology and pathway analysis was performed using Fisher’s exact test followed by a Benjamini-Hochberg correction.

## Data availability

RNA sequencing data will be available via GEO.

## Acknowledgments

AF was supported by the DFG (Deutsche Forschungsgemeinschaft) priority program 1738, SFB1286 and GRK2824;The German Federal Ministry of Science and Education (BMBF) via the ERA-NET Neuron project EPINEURODEVO; The EU Joint Programme-Neurodegenerative Diseases (JPND) – EPI-3E; Germany’s Excellence Strategy - EXC 2067/1 390729940. FS was supported by the GoBIO project miRassay (16LW0055) by the German Federal Ministry of Science and Education

## Author contribution

SS planned and conducted the majority of experiments and wrote the manuscript. MSS performed nuclei sorting MSS SB ALS and FS orchestrated RNA sequencing and DMK and TP helped with bioinformatic analysis. A.F. conceptualized the project, supervised progress and wrote the manuscript. FS and AF provide funding. SS is part of the IMPRS graduate school for Genome Science.

## Competing interests

The authors declare no conflict of interests.

## Supplemental figures

**Fig S1.**
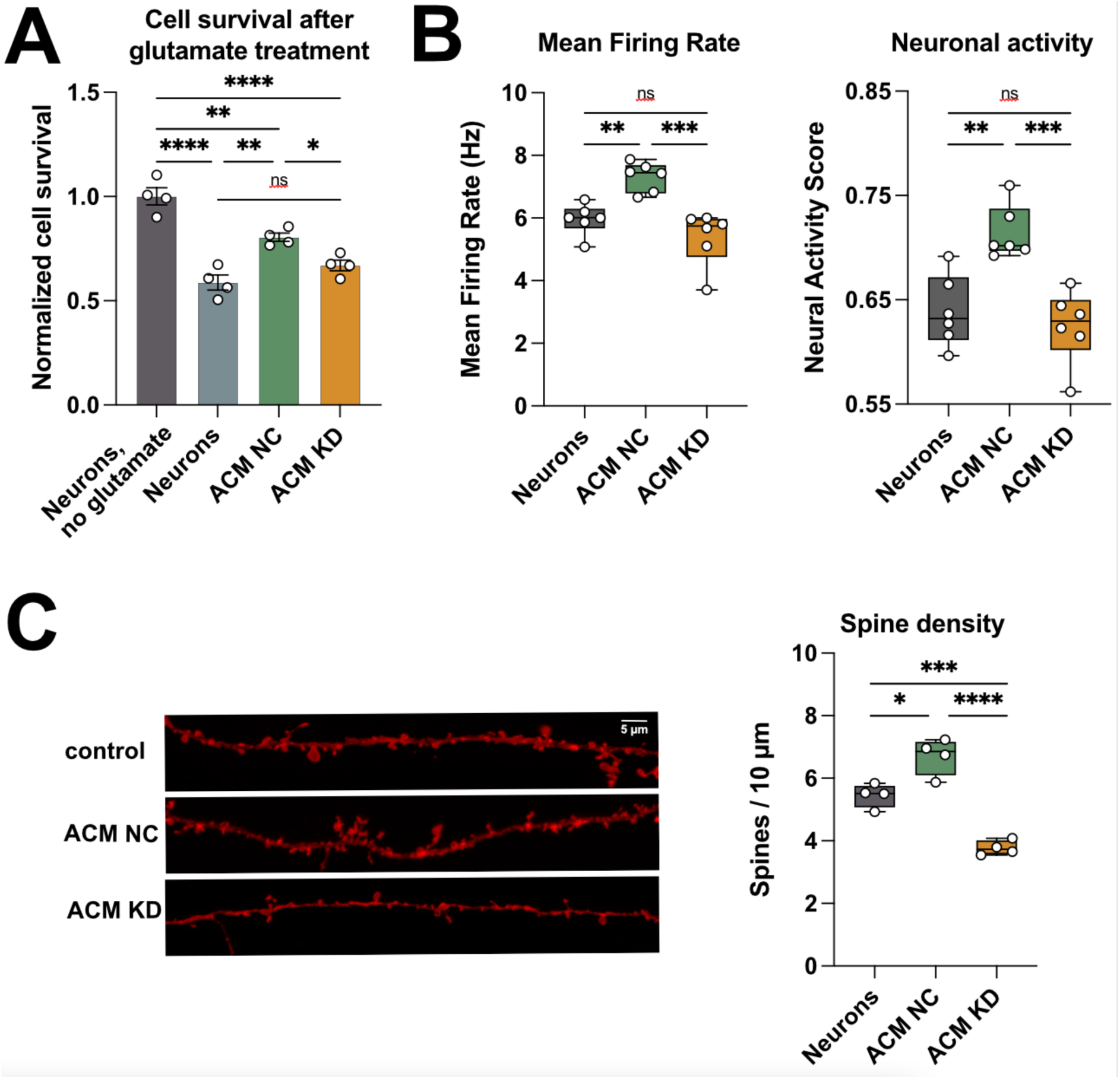
Knock down of *3222401L13Rik* in astrocytes impairs neuronal plasticity. The astrocyte-neuron co-culturing experiments shown in Fig. 4E & F are not feasible in the context of MEA assays, since co-culturing is not possible when cells are grown on MEA plates. Thus, we used a different approach to test electrophysiological properties in neurons upon astrocytic knock down of *3222401L13Rik*. To this end we treated primary astrocytes with GapmeRs targeting *3222401L13Rik* (KD), using or GapmerRs that do not bind any transcript as negative control (NC). Subsequently we harvested the media from these cells (astrocyte conditioned media, ACM) and added this to neurons followed by functional analysis. More specifically in all experiments we added to neuronal cell cultures either as control pure NB+ (media control) or ACM from primary astrocytes treated with NC GapmerRs (ACM NC) or primary astrocytes in which *3222401L13Rik* was knocked down (ACM KD). **(A)** Survival of neurons after treatment with 100 µM glutamate cultured with control medium (media control), with ACM NC or ACM KD media (One-way ANOVA; *p < 0.05, **p < 0.01, ****p < 0.0001, ns = not significant). (**B)** Left panel: mean firing rate of neurons cultured with normal medium or with ACM from NC or KD astrocytes (One-way ANOVA; **p < 0.01, ***p < 0.001, ns = not significant). Right panel: Bar plot showing the neural Activity Score obtained from MEA recordings of neurons cultured with normal medium or with ACM from NC or KD astrocytes (One-way ANOVA; **p < 0.01, ***p < 0.001, ns = not significant). (**C)** Left panel: Representative images of dendrite and spine labeling of neurons cultured with normal medium or with ACM from NC or KD astrocytes. Right panel: Quantification of spines shown in the left panel (One-way ANOVA; *p < 0.05, ***p < 0.001, ****p < 0.0001). Data are depicted as mean ± standard error. NC: negative control, KD: knockdown.

## Notes

### Competing Interest Statement

The authors have declared no competing interest.

